# N-terminal targeting sequences and coding sequences act in concert to determine the localization and trafficking pathway of apicoplast proteins in *Toxoplasma gondii*

**DOI:** 10.1101/2023.06.20.545694

**Authors:** Sofia Anjum, Aparna Prasad, Pragati Mastud, Swati Patankar

**Author notes:** Present address: University of Antwerp, Antwerp, Belgium (Aparna Prasad). Present address: Cactus Communications, Mumbai, India (Pragati Mastud). Contributed equally (Sofia Anjum, Aparna Prasad).

## Abstract

*Toxoplasma gondii* has a relict plastid, the apicoplast, to which proteins are targeted after synthesis in the cytosol. Proteins exclusively found in the apicoplast use a Golgi-independent route for trafficking, while dually targeted proteins found in both the apicoplast and the mitochondrion use a Golgi-dependent route. For apicoplast targeting, N-terminal signal sequences have been shown to direct the localization of different reporters. In this study, we use chimeric proteins to dissect out the roles of N-terminal sequences and coding sequences in apicoplast localization and the choice of the trafficking route. We show that when the N-termini of a dually targeted protein, *Tg*TPx1/2, or of an apicoplast protein, *Tg*ACP, are fused with the reporter protein, enhanced Green Fluorescent Protein (eGFP) or endogenous proteins, *Tg*SOD2, *Tg*SOD3, *Tg*ACP or *Tg*TPx1/2, the chimeric proteins exhibit flexibility in apicoplast targeting depending on the coding sequences. Further, the chimeras that are localized to the apicoplast use different trafficking pathways depending on the combination of the N-terminal signals and the coding sequences. This report shows, for the first time, that in addition to the N-terminal signal sequences, targeting and trafficking signals also reside within the coding sequences of apicoplast proteins.

## Introduction

The phylum Apicomplexa includes the obligate intracellular parasites *Toxoplasma gondii* and *Plasmodium* species that are clinically relevant. Most of these parasites, including *T. gondii*, contain a relict plastid, the apicoplast (McFadden *et al*., 1996), which though non-photosynthetic, harbors several essential metabolic pathways and is indispensable for parasite growth and survival (Jomaa *et al*., 1999; Ralph *et al*., 2004; Mazumdar *et al*., 2006). The apicoplast was procured by secondary endosymbiosis, where an autotrophic alga containing a primary plastid was engulfed by another eukaryote (Bodył, 1999; Parsons *et al*., 2007). Acquisition of the apicoplast was followed by gene transfer to the host nucleus, an event that conferred host control over the endosymbiont and confined the free-living eukaryote into a semi-autonomous organelle (reviewed in Waller & McFadden, 2005). Due to the transfer of genes to the nucleus, the apicoplast relies on the cell for translation of the newly acquired genes; therefore, apicomplexan parasites must use the cellular trafficking machinery to send nuclear-encoded proteins to the apicoplast.

Trafficking of nuclear-encoded proteins to different destinations is achieved by the presence of specific signals often present at the N-terminus of the protein sequence. For example, N-terminal signal peptides (SP) direct secretory proteins through the endoplasmic reticulum (ER) and eventually to the secretory pathway (von Heijne, 1990) and an N-terminal mitochondrial targeting sequence (MTS) direct proteins to the mitochondrion (Roise and Schatz, 1988). For most apicoplast proteins, an N-terminal bipartite signal, an SP followed by a transit peptide (TP), serves as a specific tag for organellar localization (Waller *et al*., 1998). The SP directs proteins into the ER, where it gets cleaved, exposing the TP, which then directs proteins to the apicoplast and is removed once inside the organelle, releasing the mature protein (Waller *et al*., 1998, 2000; Ralph, 2004; Parsons *et al*., 2007). Hence, the ER is the first step in apicoplast protein targeting. Interestingly, proteins from the ER can take two routes to the apicoplast, i.e., the classical secretory pathway via the Golgi or a Golgi-independent pathway (DeRocher *et al*., 2005; Prasad, Mastud and Patankar, 2021).

Even though specific signals ensure localization to a particular compartment, there are reports of a single protein localizing in multiple organelles (Chew *et al*., 2003; Pino *et al*., 2007; Baudisch *et al*., 2014). In *T. gondii*, thioredoxin peroxidase, superoxide dismutase, and aconitase (*Tg*TPx1/2, *Tg*SOD2, and *Tg*ACN) dually localize to the apicoplast and the mitochondrion (Pino *et al*., 2007). A detailed study of the N terminal signal sequences of *Tg*TPx1/2 revealed that this dually localized protein has a single targeting sequence for both the apicoplast and the mitochondrion, as opposed to two distinct sequences (Mastud and Patankar, 2019).

Organelle-targeting signal sequences have been used to deliver reporters and cellular proteins to particular organelles (Huttly, 2009). In *T. gondii,* enhanced Green Fluorescent Protein (eGFP), among others, is a commonly employed fluorescent reporter for studying various aspects of parasite biology (Gubbels and Striepen, 2004; Striepen and Soldati, 2007; Hanig, Entzeroth and Kurth, 2012). The N-terminal targeting signals of exclusively apicoplast-targeted proteins are capable of localizing GFP to the apicoplast (Waller *et al*., 1998; DeRocher *et al*., 2005), however, the N-terminal sequences of *Tg*SOD2 (a protein dually localized to the apicoplast and mitochondrion) were unable to target GFP to the apicoplast, with the fusion protein showing only mitochondrial localization (Brydges and Carruthers, 2003; Pino *et al*., 2007). This report extends these published observations to another dually localized protein, *Tg*TPx1/2. Like *Tg*SOD2, a loss of apicoplast localization was observed when enhanced GFP (eGFP) was fused with the N-terminal signal sequences of *Tg*TPx1/2. However, apicoplast localization was restored when eGFP was replaced with endogenous proteins of *T. gondii*.

For the first time, we investigate whether the choice of the reporter protein can also affect the choice of the trafficking pathway to the apicoplast. We ask whether the N-terminal targeting sequences alone are sufficient to direct a reporter protein through the trafficking route that is taken by the native protein. We show that the N-terminal sequence of *Tg*TPx1/2 (a dually targeted protein that takes the Golgi-dependent route to the apicoplast) can drive apicoplast localization of *Tg*SOD3, a mitochondrial protein, through the same route. Similarly, the N-terminal sequence of *Tg*ACP (an apicoplast protein that takes the Golgi-independent route), drives the localization of *Tg*TPx1/2 through the same route. Surprisingly, when the N-terminal sequence of *Tg*TPx1/2 is fused with the coding sequences of *Tg*ACP, and also when the N-terminal sequence of *Tg*ACP is fused with *Tg*SOD2, the expected trafficking route (based on the N-terminal sequence) switches to the other pathway. Therefore, the choice of the trafficking pathway of chimeric proteins depends on specific combinations of signal sequences and coding sequences, indicating, for the first time, that apicoplast targeting motifs are present throughout the entire protein coding sequence. This work has implications for the field of apicoplast biology, which has been largely dominated by studies of apicoplast proteins where the N-terminal sequences are sufficient to drive proteins to their destinations.

## Materials and Methods

### Plasmid construction of *T. gondii* genes in expression vectors

For constructing C-terminal eGFP fusion proteins of *Tg*TPx1/2 (ToxoDB gene ID: TGGT1_266120), pCTG-TPx1/2-HA (Mastud and Patankar, 2019) was used as a template for amplifying the full-length protein (TPx1/2_FL_) and the N-terminus encoding the 1-150 amino acids (TPx1/2_N_). These DNA sequences were cloned between the *NheI* and *AvrII* sites of pCTG-eGFP (van Dooren et al., 2008), generating pCTG-TPx1/2FL-eGFP and pCTG-TPx1/2N-eGFP, respectively. The TPx1/2(R24A)N-eGFP plasmid was generated by site-directed mutagenesis, replacing the arginine at the position 24 with alanine.

For the generation of pCTG-TPx1/2_N_-SOD2-HA and pCTG-TPx1/2_N_-SOD3-HA plasmids, first, the N-terminus of TPx1/2_N_(1-150 amino acids) was amplified from pCTG-TPx1/2-HA and cloned between *MfeI* and *NdeI* sites in the pCTG-HA vector. The coding sequences of *Tg*SOD2 (ToxoDB gene ID: TGGT1_316330) from 92-287 amino acids and *Tg*SOD3 (ToxoDB gene ID: TGGT1_316190) from 57-258 amino acids were amplified from pCTG-SOD2-HA and pCTG-SOD3-HA (Prasad, Mastud and Patankar, 2021) and inserted downstream of the TPx1/2_N_ in the *NdeI* site of the pCTG-TPx1/2_L_-HA vector. For generating TPx1/2_N_-SOD2-HA-HDEL and TPx1/2_N_-SOD3-HA-HDEL constructs, the respective genes were amplified using a common forward primer and a reverse primer containing an HDEL sequence and were cloned in the pCTG-HA vector.

For the construction of ACP_N_-eGFP, the N-terminal 104 amino acids of ACP (ToxoDB gene ID: TGGT1_264080) were amplified from cDNA and cloned between the *MfeI* and *NdeI* sites of pCTG-HA vector. This ACP_N_ (1-104 amino acids) was then amplified and cloned upstream of eGFP between the *NheI* and *AvrII* sites of the pCTG-eGFP vector. The HA and HDEL constructs of TPx1/2_N_-ACP were constructed by replacing the coding sequence of SOD2 in the *NdeI* site from the HA, and HA-HDEL tagged pCTG-TPx1/2_N_-SOD2 vector with the coding sequence of ACP (105-183 amino acids) amplified from cDNA.

For the generation of HA and HA-HDEL constructs of ACP_N_-TPx1/2 and ACP_N_-SOD2, first, pCTG-ACP_N_-HA and pCTG-ACP_N_-HA-HDEL were constructed by amplifying the first 104 amino acids of ACP_L_ from pCTG-ACP_N_-eGFP and replacing it in the location of the ATrx1 gene between the *MfeI* and *NdeI* sites in the pCTG-ATrx1-HA and pCTG-ATrx1-HA-HDEL vectors (Prasad, Mastud and Patankar, 2021). The coding sequences of *Tg*TPx1/2 (151-333 amino acids) and *Tg*SOD2 (92-287 amino acids) were then amplified from pCTG-TPx1/2-HA and pCTG-SOD2-HA and inserted downstream of ACP_N_ in the *NdeI* site of pCTG-ACP_N_-HA and pCTG-ACP_N_-HA-HDEL vector. All the clones were confirmed by sequencing and found to be devoid of mutations, except pCTG-ACP_N_-SOD2-HA and pCTG-ACP_N_-SOD2-HA-HDEL where a leucine was mutated to isoleucine at position 239, which might have been generated during PCR amplification. Since leucine and isoleucine are both branched chain amino acids this mutation should not result in a different phenotype. The primers used are listed in Table 1.

**Table 1:**
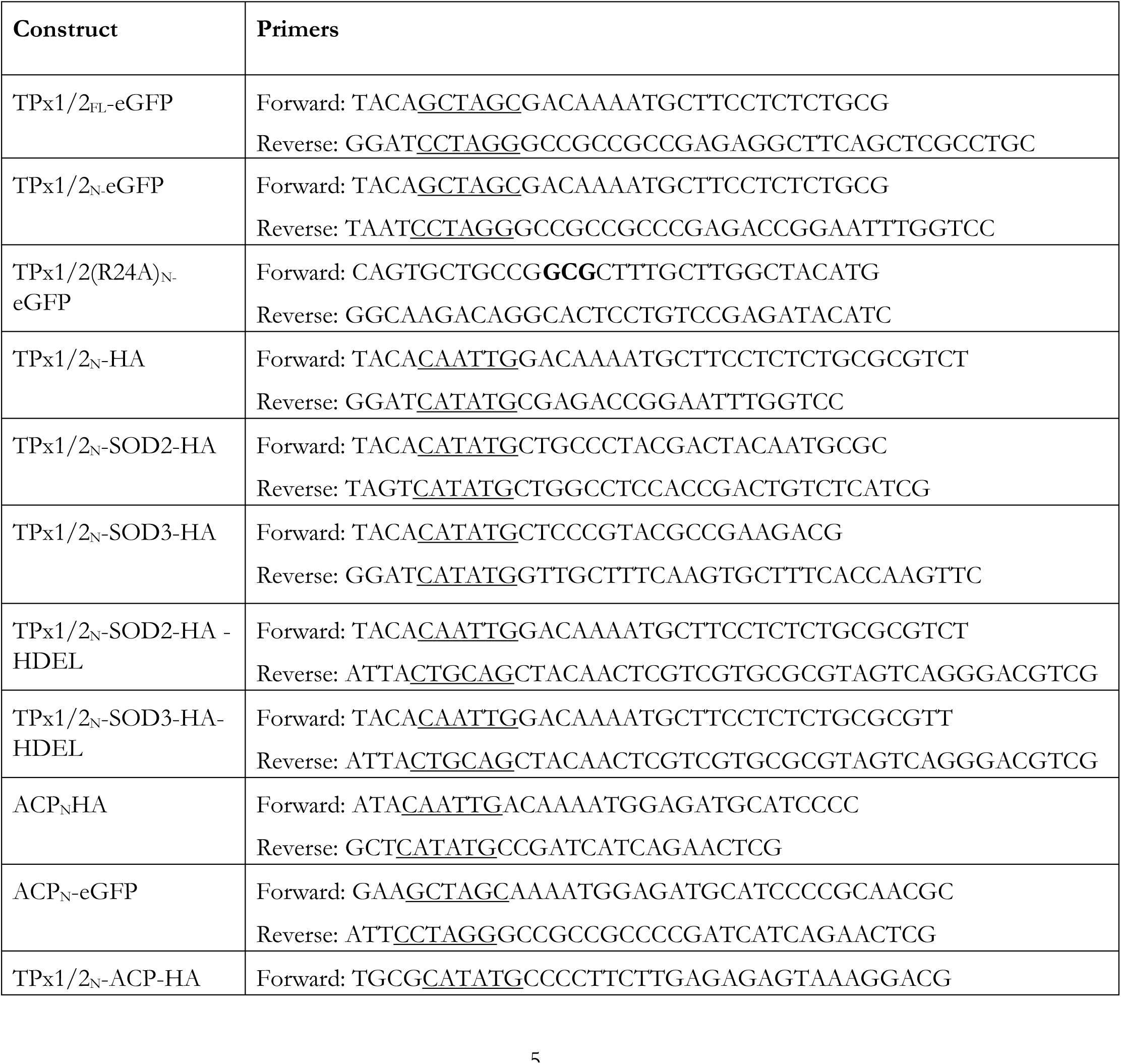
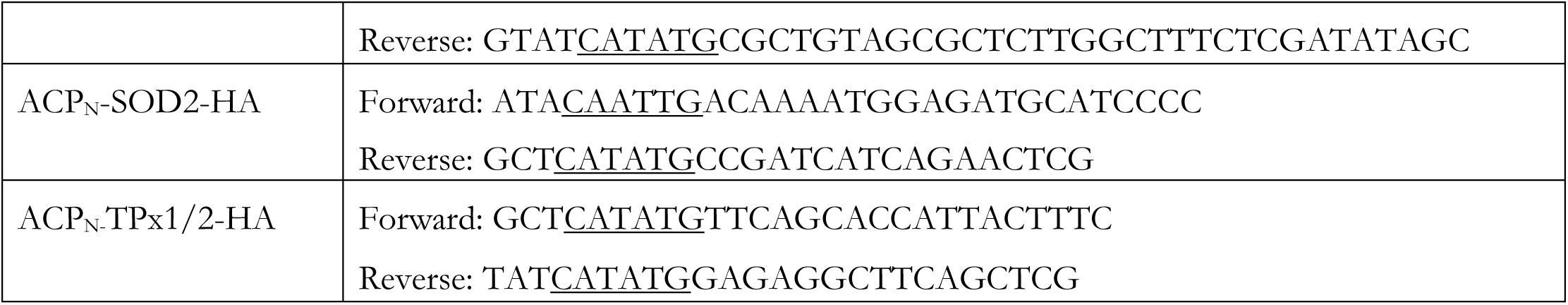
List of primers used in this study. All the primers are shown in the 5′ to 3′ direction with the restriction sites underlined, and the desired mutations shown in bold.

### Parasite maintenance and transient transfections

*T. gondii* wild-type RH strain were cultured and maintained in primary human foreskin fibroblast (HFF) cells as described previously (Striepen and Soldati, 2007). For transient transfection of tachyzoites by a single plasmid, parasites were harvested and resuspended in a reaction mixture of 400 µl of incomplete DMEM (without serum and gentamicin) with 50 µg DNA of plasmid. For transient co-transfection, extracellular parasites were resuspended in Cytomix buffer containing 120 mM KCl (Merck), 0.15 mM CaCl (Merck), 10 mM K_2_HPO_4_ (SRL Chemical), 10 mM KH_2_PO_4_ (Sisco Research Laboratories) pH 7.6, 25 mM HEPES pH 7.6 (Sigma-Aldrich), 2 mM EGTA (Merck), and 5 mM MgCl_2_ (Merck) (Striepen and Soldati, 2007) with 35 μg of DNA of each plasmid. These reaction mixtures were then added in a 2 cm Bio-Rad GenePulser electroporation cuvette and electroporated at 1.5 kV, 50 Ω, and 25 µF using a Bio-Rad GenePulser Xcell system. Transfected parasites were grown in HFF cells for 24 hours, followed by immunostaining or western blot.

### Immunostaining and microscopy

For immunostaining, transiently transfected parasites were grown in chamber slides (SPL Life Science Co.). After 24 hours, intracellular parasites were fixed in 4% formaldehyde (Merck) and 0.0075% glutaraldehyde in 1X PBS for 30 minutes. After fixing, parasites were washed twice with 1X PBS and were permeabilized by treating with 0.25% Triton X-100 (Sigma-Aldrich) for 10 minutes. Blocking was performed with 3% BSA (GeNei^TM^) for one hour. After blocking, parasites were incubated with primary antibody, 1:1000 anti-HA rabbit monoclonal antibody (C29F4; Cell Signaling Technology), 1:1000 anti-HA mouse monoclonal antibody (Clone HA-7, mouse IgG1 Isotype; Sigma-Aldrich), or 1:1000 anti-ACP antibodies (a kind gift from Prof. Dhanasekaran Shanmugam, NCL, Pune) for 2 hours at room temperature followed by washing with 1X PBS. After three washes, parasites were incubated with appropriate secondary antibody for 1.5 hours at room temperature (anti-rabbit Alexa 488 (1:400), anti-rabbit 568 (1:400) and anti-mouse Alexa 594 (1:250) from Invitrogen^TM)^. Parasites were washed thrice in 1X PBS and mounted in ProLong Diamond Antifade mountant (Thermo Fisher Scientific). MitoTracker^®^ red CMXRos (300 nM) from Invitrogen^TM^ was used for staining mitochondria as described previously (Salunke *et al*., 2018)

Confocal images were acquired by a Zeiss LSM 780 NLO confocal microscope using a Plan-Apochromat 100X, 1.4 NA objective. Optical sections along the Z-axis were captured and projected at maximum intensity.

### Image processing and analysis

To investigate the colocalization of proteins with organelle markers, we microscopically observed 40 vacuoles of transfected parasites, and ∼10 parasite vacuoles were imaged randomly for documentation and analyzed for colocalization by either Imaris software or Image J. For comparative analysis of the localization of proteins with and without the C-terminal HDEL sequence with the apicoplast marker, at least 40 parasite vacuoles were randomly imaged using the same settings. We used unprocessed images of parasite vacuoles to quantitatively analyze the colocalization of the protein of interest with the apicoplast marker by Pearson’s correlation coefficient (PCC) with Image J software. A region of interest was selected within a parasite, covering the entire volume of the apicoplast marker, and an automatic threshold was applied. A dot plot of PCC values showing the median with a 95% Confidence Interval was then plotted using GraphPad Prism version 8.0.1(244) for Windows, GraphPad Software, La Jolla, California, USA. The statistical significance of the change in the PCC values of proteins with and without HDEL with apicoplast marker was analyzed using the Mann-Whitney test. Representative images of parasites expressing fusion proteins were later processed for brightness and contrast in ImageJ; however, no non-linear adjustments were made, such as changes in the gamma settings. All the experiments were performed in duplicates.

### Comparison between different approaches for studying apicoplast localization with different apicoplast markers

Due to logistics issues regarding antibodies recognizing *Tg*ACP, the apicoplast marker, we were forced to use ACP_N_-eGFP as the apicoplast marker midway through the study. First, we checked the co-transfection efficiency of ACP_N_-eGFP with ACP_N_-SOD2-HA. The co-transfection efficiency ranged between ∼30-40% of the transfected parasites, showing that doubly transfected parasites could be easily detected for further immunofluorescence analysis. To compare the co-transfection strategy to the experiments that used anti-*Tg*ACP antibodies for staining the apicoplast, we co-transfected plasmids expressing *Tg*ACP_N_-eGFP with all the constructs. No significant difference in the immunofluorescence or the median PCC values was observed between the two strategies; therefore, the co-transfection strategy was deemed suitable for the remaining experiments.

### Western blotting

For stable transfectants, ∼10 million parasites were lysed in SDS-PAGE gel loading buffer (1X) and loaded on a 15% SDS-PAGE gel. The proteins were transferred onto a PVDF membrane at 35mA for 16 hours. The membrane was blocked with 3% Skimmed milk (Himedia) in PBST for 1 hour 30 mins and incubated with primary antibody for 2 hours at a dilution of 1:1000 in 3% BSA-PBST. Primary antibodies used anti-HA Rabbit monoclonal antibody (C29F4; Cell Signaling Technology), anti-HA Mouse monoclonal antibody (Clone HA-7, mouse IgG1 Isotype; Sigma-Aldrich). The membrane was washed thrice in PBST and incubated with HRP-conjugated secondary antibody (anti-rabbit and anti-mouse; GeNei^TM^) for 1 hour at a dilution of 1:2000 in 3% BSA-PBST. The blot was developed as per instructions for chemiluminescence using Clarity Western ECL Substrate (1705061; Bio-Rad). The relative density was measured using Image J software.

## Results

Most of the apicoplast proteins reported in the literature, N-terminal signal sequences are sufficient to direct proteins to their destination (Waller *et al*., 1998; DeRocher *et al*., 2000; He *et al*., 2001). However, for *Tg*SOD2, a protein dually localized to both the apicoplast and the mitochondrion, the N-terminus drives different reporter proteins to different organelles (Pino *et al*., 2007); the trafficking routes taken by these proteins to the apicoplast has not been tested. To understand the inherent nature of targeting signals in apicoplast proteins, we made several chimeric constructs and analyzed their effects on both localization and trafficking routes to the apicoplast.

### The N-terminus of *Tg*TPx1/2 drives eGFP to the mitochondrion, with a loss of apicoplast targeting

*Tg*TPx1/2 is a dually localized protein, which localizes to both the apicoplast and the mitochondrion, when tagged with small epitope tags such as hemagglutinin (HA) (Pino *et al*., 2007). Due to their advantages during live-cell imaging, reporter proteins like GFP have been used in *T. gondii* for visualizing organelles (Waller *et al*., 1998; Brydges and Carruthers, 2003). Therefore, we assessed the localization of an in-frame fusion of *Tg*TPx1/2_FL_ with enhanced GFP (eGFP) at the C-terminus and found that this protein showed a distinct mitochondrial localization, with no colocalization with the apicoplast marker (Fig. 1A). This observation suggested that the eGFP reporter protein interferes with the apicoplast targeting of *Tg*TPx1/2.

**Figure 1:**
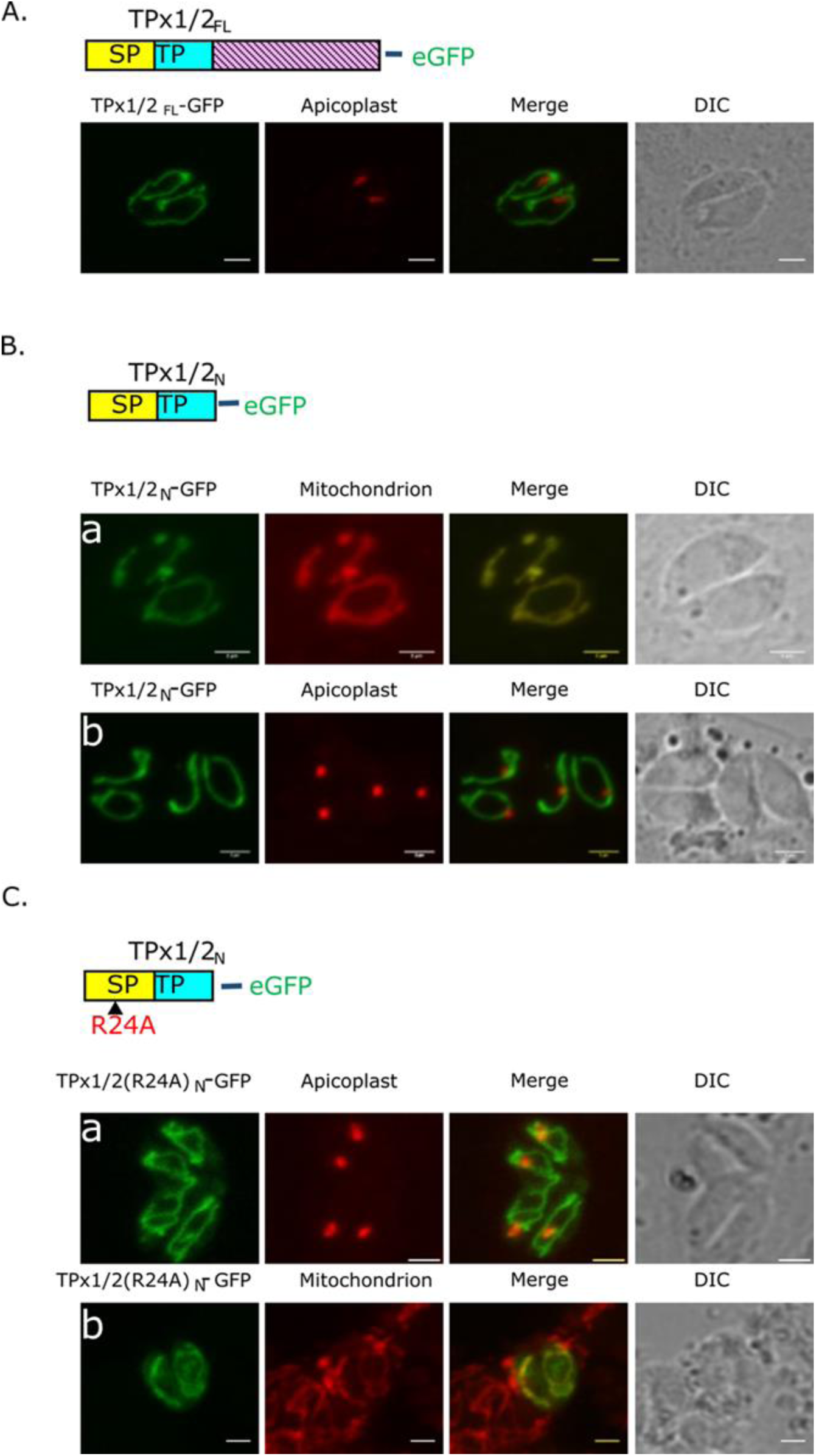
A GFP reporter results in the localization of *Tg*TPx1/2 to the mitochondrion but not the apicoplast. Microscopic images of parasites transiently expressing varying lengths of *Tg*TPx1/2 protein (green) fused with eGFP. The N-terminus of *Tg*TPx1/2, consisting of the signal peptide (SP) and transit peptide (TP), is shown in yellow and blue, respectively, while the mature protein is shown in purple **A)** The full-length protein of *Tg*TPx1/2 fused with eGFP at its C-terminus [*Tg*TPx1/2_FL_-eGFP] (green) with the apicoplast marker *Tg*ACP (red), **B)** The N-terminus of *Tg*TPx1/2_N_ (SP + TP) fused with eGFP at its C-terminus [*Tg*TPx1/2_N_-eGFP] (green), with the apicoplast marker *Tg*ACP in red (panel a) and the mitochondrion marker, SP-TP-SOD2-DsRed in red (panel b) **C)** The N-terminus of *Tg*TPx1/2 with an (R24A) mutation fused with eGFP [*Tg*TPx1/2(R24A)_N_-eGFP] (green), with the apicoplast marker *Tg*ACP in red (panel a) and the mitochondrion marker, Mitotracker in red (panel b). The apicoplast marker was detected using anti-*Tg*ACP antibodies. Scale bar = 2 µm.

Organellar targeting signals for apicoplast and mitochondrial import are present at the N-terminus of the protein (Sheiner and Soldati-Favre, 2008). For *Tg*TPx1/2, it was previously demonstrated that the first 50 amino acids of the protein fused to eGFP showed faint apicoplast and mitochondrial localization (Mastud and Patankar, 2019). Interestingly, a Western blot of this protein revealed that the signal cleavage sites for organellar uptake are present beyond the first 50 amino acids of the protein (Mastud and Patankar, 2019). An analysis of *Tg*TPx1/2 by the Simple Modular Architecture Research Tool (SMART) Version 9 (Letunic, Doerks and Bork, 2015; Letunic and Bork, 2018) predicted a glutathione peroxidase domain starting at amino acid 151. These data suggest that amino acids from 1-150 of *Tg*TPx1/2 should possess signals for targeting to both the apicoplast and the mitochondrion. To test this, we generated a chimeric construct encoding the first 150 amino acids of *Tg*TPx1/2 fused to eGFP (Fig. 1B) and observed that parasites expressing *Tg*TPx1/2_N_-eGFP showed colocalization with the mitochondrial marker SP-TP-SOD2-DsRed (Pino *et al*., 2007) but not with the apicoplast marker acyl carrier protein, *Tg*ACP (median PCC value 0.07 (Fig. 1B). Charged amino acid mutants in the signal peptide region *Tg*TPx1/2(R24A) completely shift the localization of *Tg*TPx1/2 to the apicoplast (Mastud and Patankar, 2019). We generated an R24A mutant of *Tg*TPx1/2_N_-eGFP and expected it to localize exclusively to the apicoplast. Contrary to expectations (Fig. 1C), *Tg*TPx1/2(R24A)_N_-eGFP localized to the mitochondrion and only showed a faint signal near the apicoplast (median PCC value 0.10).

As the full-length protein, the first 150 amino acids of *Tg*TPx1/2 and an R24A mutant of *Tg*TPx1/2_N_ when fused to eGFP were all localized to the mitochondrion, we infer that eGFP interferes with the apicoplast localization of this dually targeted protein. This result is consistent with data seen for another dually targeted protein, *Tg*SOD2 (Brydges and Carruthers, 2003; Pino *et al*., 2007), and in complete contrast to data for proteins that are localized only to the apicoplast (Waller *et al*., 1998; DeRocher *et al*., 2000; Wu *et al*., 2015)

### The N-terminus of *Tg*TPx1/2 targets endogenous proteins to the apicoplast and weakly to the mitochondrion

As the heterologous reporter protein, eGFP affects the localization of *Tg*TPx1/2 to the apicoplast, we employed an endogenous protein of *T. gondii*, superoxide dismutase 3 (*Tg*SOD3), with a C-terminal HA tag as a reporter protein. *Tg*SOD3 is a mitochondrial protein containing a mitochondrial targeting sequence from residues 1-56 (Brydges and Carruthers, 2003). The first 150 amino acids of *Tg*TPx1/2 were fused to *Tg*SOD3 (amino acids 57-258), devoid of its targeting signals. In contrast to *Tg*TPx1/2_N_-eGFP, *Tg*TPx1/2_N_-SOD3 was targeted to the apicoplast in the entire population of transiently transfected parasites (Fig. 2A, 2C), while ∼68% of parasites showed both apicoplast and the mitochondrion signals (Fig. 2A).

**Figure 2:**
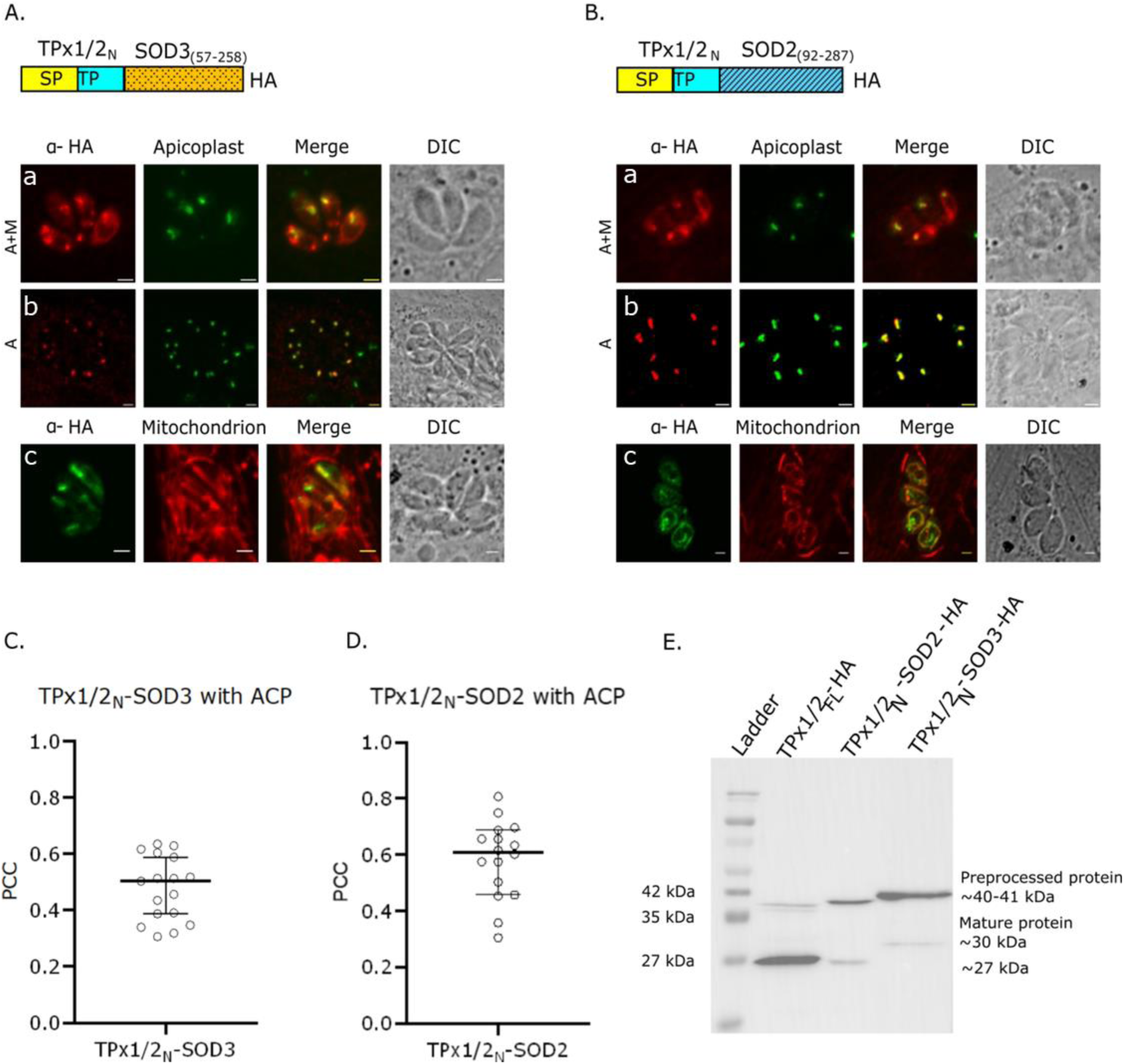
**The N-terminus of the dually targeted protein *Tg*TPx1/2 can target endogenous proteins to the apicoplast and the mitochondrion**. Microscopic images of parasites expressing HA-tagged fusion proteins consisting of the N-terminus of *Tg*TPx1/2 (SP + TP) fused in-frame with the coding sequences of endogenous *T. gondii* proteins (*Tg*SOD2 and *Tg*SOD3) are shown in red. These chimeric proteins show two phenotypes: only Apicoplast (A) and Apicoplast + Mitochondrion (A+M) **A)** *Tg*TPx1/2_N_-SOD2, **B)** *Tg*TPx1/2_N_-SOD3 with the apicoplast marker *Tg*ACP in green (panel a and b) and the mitochondrial marker MitoTracker in green (panel c), **(C, D)** The graphs showing median PCC value with 95% confidence interval of *Tg*TPx1/2_N_-SOD2 (median PCC value 0.5) and *Tg*TPx1/2_N_-SOD3 (median PCC value 0.6) with apicoplast marker *Tg*ACP, **E)** Western blot analysis using anti-HA antibodies of the cell lysate of parasites transiently transfected with *Tg*TPx1/2_FL_-HA, *Tg*TPx1/2_N_-SOD2, and *Tg*TPx1/2_N_-SOD3. Signal peptide (SP) is shown in yellow, transit peptide (TP) is in blue, the mature protein of *Tg*SOD3 is shown with a pattern in yellow, and the mature protein of *Tg*SOD2 is shown with a pattern in blue. The apicoplast marker was detected using anti-*Tg*ACP antibodies. Scale bar = 2 µm.

When the N-terminus of *Tg*TPx1/2 was fused to the coding sequences of another dually targeted protein, *Tg*SOD2 (92-287), the fusion protein *Tg*TPx1/2_N_-SOD2 was targeted to the apicoplast in the entire population (Fig. 2B, 2D). This fusion protein was present in the apicoplast and the mitochondrion in ∼63% of the population (Fig. 2B). Similar to the observations for *Tg*SOD3, the mitochondrial signal was weak (Fig. 2B). Therefore, unlike eGFP, both *Tg*TPx1/2_N_-SOD3 and *Tg*TPx1/2_N_-SOD2 show robust apicoplast targeting and weak mitochondrial targeting. This result is different from another dually targeted protein, *Tg*SOD2. The N-terminus of *Tg*SOD2, when fused to eGFP, is targeted only to the mitochondrion, but when fused to the coding sequence of *Tg*SOD3, dual targeting is restored (Pino *et al*., 2007).

To confirm that the chimeric proteins are correctly processed, we performed western blots of the transiently transfected parasites using antibodies against the HA tag (Fig. 2E). The wild type *Tg*TPx1/2 and the chimeric constructs *Tg*TPx1/2_N-_SOD2, *Tg*TPx1/2_N-_SOD3 showed pre-processed protein and a mature protein of the expected sizes (∼40-41 kDa and ∼27-30 kDa, respectively), indicating that the N-terminal targeting signals have the appropriate sequences for cleavage in the apicoplast and mitochondrion.

### The coding sequences of apicoplast proteins contain signals that drive them through the Golgi-independent pathway

Since the presence of endogenous proteins restored the apicoplast localization of fusion proteins having N-terminal signal sequences of *Tg*TPx1/2, we next checked the choice of pathway of these fusion proteins to the apicoplast. Dually targeted proteins are known to take a Golgi-dependent pathway to the apicoplast, while exclusively apicoplast-localized proteins are targeted independently of the Golgi (Prasad, Mastud and Patankar, 2021). We hypothesized that if the N-terminal 150 amino acids of *Tg*TPx1/2 with endogenous proteins can reach the apicoplast, the fusion protein should contain all the signals for trafficking through a Golgi-dependent pathway.

To test this hypothesis, an HDEL sequence was added at the C-terminal to *Tg*TPx1/2_N_-SOD3-HA and *Tg*TPx1/2_N_-SOD2-HA (Fig. 3A and 3B). In *T. gondii*, the HDEL motif serves as a retrieval signal for ER proteins that have escaped to the Golgi apparatus by bulk flow (Hager *et al*., 1999), and if a protein employs a Golgi-dependent route for trafficking, the addition of the HDEL motif results in a reduced localization of the protein to the destined compartment and an increase in ER retention (DeRocher *et al*., 2000; Prasad, Mastud and Patankar, 2021). Since *Tg*TPx1/2_N_-SOD3-HA and *Tg*TPx1/2_N_-SOD2-HA are dually localized to the apicoplast and the mitochondrion, it was difficult to distinguish the increase in the perinuclear staining from the mitochondrion signal of the HDEL constructs of these chimeric proteins. Also, previously, it was shown for *Tg*SOD2, a dually targeted protein that employs an ER-Golgi route for apicoplast trafficking, the addition of the HDEL sequence resulted in a reduced apicoplast localization without any increase in ER retention (Prasad, Mastud and Patankar, 2021). Therefore, ER retention was not assayed; instead, a quantitative analysis of the reduced signal of the HDEL-containing protein in the apicoplast was measured. Reduced signal of chimeric proteins with apicoplast marker would indicate that the protein traverses through the Golgi. The Pearson Correlation coefficient was calculated for proteins with and without the HDEL sequence with the apicoplast marker, and a comparative analysis was done between the population of the two constructs. No reduction in the apicoplast localization of HDEL-containing proteins would suggest that the protein employs a Golgi-independent route. This strategy has been used previously to understand the trafficking pathway of apicoplast proteins in *T. gondii* (Prasad, Mastud and Patankar, 2021).

**Figure 3:**
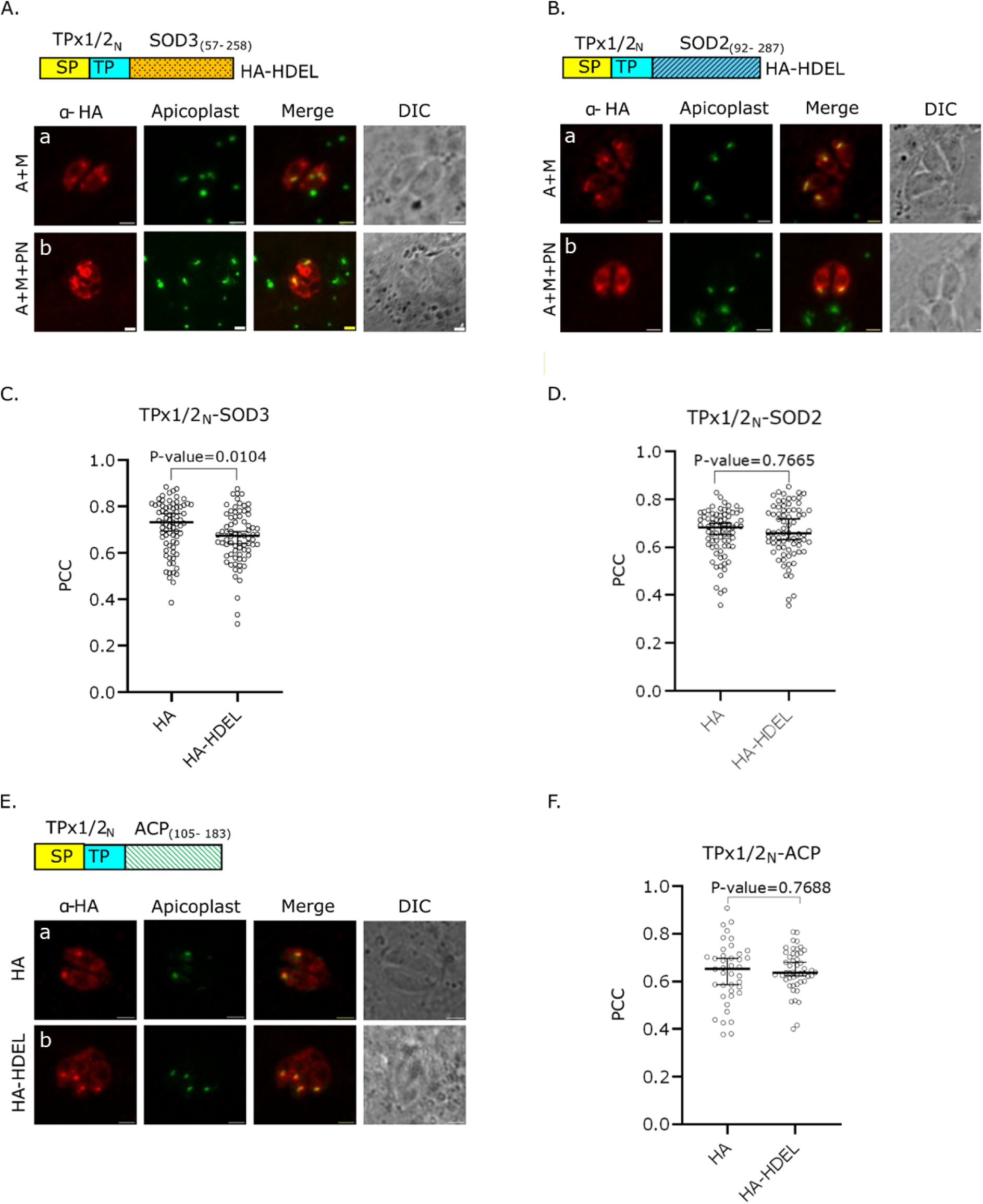
Examining the trafficking pathway of *Tg*TPx1/2 fused with endogenous *T. gondii* proteins. Microscopic images of parasites transiently expressing chimeric proteins with and without the HDEL sequence at their C-terminus are in red, and the apicoplast marker is in green. **A)** *Tg*TPx1/2_N_-SOD3-HA-HDEL **B)** *Tg*TPx1/2_N_-SOD2-HA-HDEL, with the apicoplast marker *Tg*ACP (green), exhibiting two phenotypes; Apicoplast and Mitochondrion (A+M) shown in panel a and Apicoplast and Mitochondrion with perinuclear staining (A+M+PN) shown in panel b **C, D)** Pooled PCC values of HA and HA-HDEL tagged proteins of *Tg*TPx1/2_N_-SOD3, *Tg*TPx1/2_N_-SOD2 with apicoplast marker, *Tg*ACP and *Tg*ACP-eGFP showing median value with 95% confidence interval respectively **E)** *Tg*TPx1/2_N_-ACP (red) with and without HDEL sequence (panel a and b respectively) with the apicoplast marker *Tg*ACP-eGFP (green). **F)** Pooled PCC values of HA and HDEL tagged proteins of *Tg*TPx1/2_N_-ACP with apicoplast marker, *Tg*ACP-eGFP showing median value with 95% confidence interval. The apicoplast marker was detected using anti-*Tg*ACP antibodies. Scale bar = 2 µm.

For *Tg*TPx1/2_N_-SOD3-HA-HDEL and *Tg*TPx1/2_N_-SOD2-HA-HDEL, two phenotypes were observed, with one showing dual localization (Fig. 3A and 3B, panel a) and another showing dual localization with an increase in the perinuclear spread (Fig. 3A and 3B, panel b). However, for *Tg*TPx1/2_N_-SOD3-HA-HDEL, a significant decrease in the PCC values (P-value: 0.0104) of the HDEL construct was observed when compared to the construct without an HDEL motif (Fig. 3C). This suggests that the N-terminus of *Tg*TPx1/2 is able to steer *Tg*SOD3 through the Golgi-dependent pathway to the apicoplast. Contrastingly, for *Tg*TPx1/2_N_-SOD2-HA, no significant difference (P-value: 0.7665) was observed in the PCC value between the HA and the HDEL constructs, indicating a Golgi-independent route for this protein (Fig. 3D). As *Tg*SOD3 is a mitochondrial protein while *Tg*SOD2 is a dually targeted protein, we created a construct with N-terminal signal sequences of *Tg*TPx1/2 fused with the coding sequence of exclusive apicoplast protein taking a Golgi-independent route, *Tg*ACP (105-183 amino acids). Similar to *Tg*TPx1/2_N_-SOD2, parasites expressing *Tg*TPx1/2_N_-ACP with or without an HDEL motif showed neither any increase in the perinuclear staining nor any significant difference between the median PCC values (P-value: 0.7688) with the apicoplast marker *Tg*ACP_N_-GFP (Figs. 3E and 3F), indicating a Golgi-independent route for apicoplast trafficking. Therefore, the N-terminal targeting sequences of *Tg*TPx1/2 can target the mature domain of *Tg*SOD3 to the apicoplast through the Golgi, while the mature domain of the apicoplast proteins *Tg*ACP and *Tg*SOD2 appear to contain signals for trafficking through the Golgi-independent route.

### The coding sequences of dually targeted proteins play a role in apicoplast trafficking through the Golgi

The exclusively apicoplast-targeted protein *Tg*ACP has an N-terminal signal sequence from 1-104 amino acids (*Tg*ACP_N_) that is sufficient to drive exogenous reporters like GFP and DsRed to the apicoplast (Waller *et al*., 1998; Pino *et al*., 2007) using a Golgi-independent route (DeRocher *et al*., 2005). We asked whether this signal sequence can drive dually targeted proteins to the apicoplast through a Golgi-independent pathway.

The N-terminal signal sequences of *Tg*ACP (1-104 amino acids) was fused with the coding sequences of the dually targeted protein *Tg*TPx1/2 (151-333 amino acids). Parasites expressing *Tg*ACP_N_-TPx1/2-HA showed apicoplast localization in 60% of the population and apicoplast localization with perinuclear staining in 40% of the population (Fig. 4, panel a), similar to previous reports with other proteins (Waller *et al*., 1998; He *et al*., 2001; van Dooren *et al*., 2002; DeRocher *et al*., 2005; Karnataki *et al*., 2009). To check the trafficking pathway of *Tg*ACP_N_-TPx1/2-HA, we added an HDEL motif to its C-terminus. Microscopic images of the HDEL construct of *Tg*ACP_N_-TPx1/2 showed no increase in the perinuclear staining, as shown in Fig. 4A (panel b). Additionally, the median PCC values of the *Tg*ACP_N_-TPx1/2-HA protein, with or without an HDEL motif, showed no significant difference with a P-value of 0.8030, indicating a Golgi-independent route (Fig. 4B).

**Figure 4:**
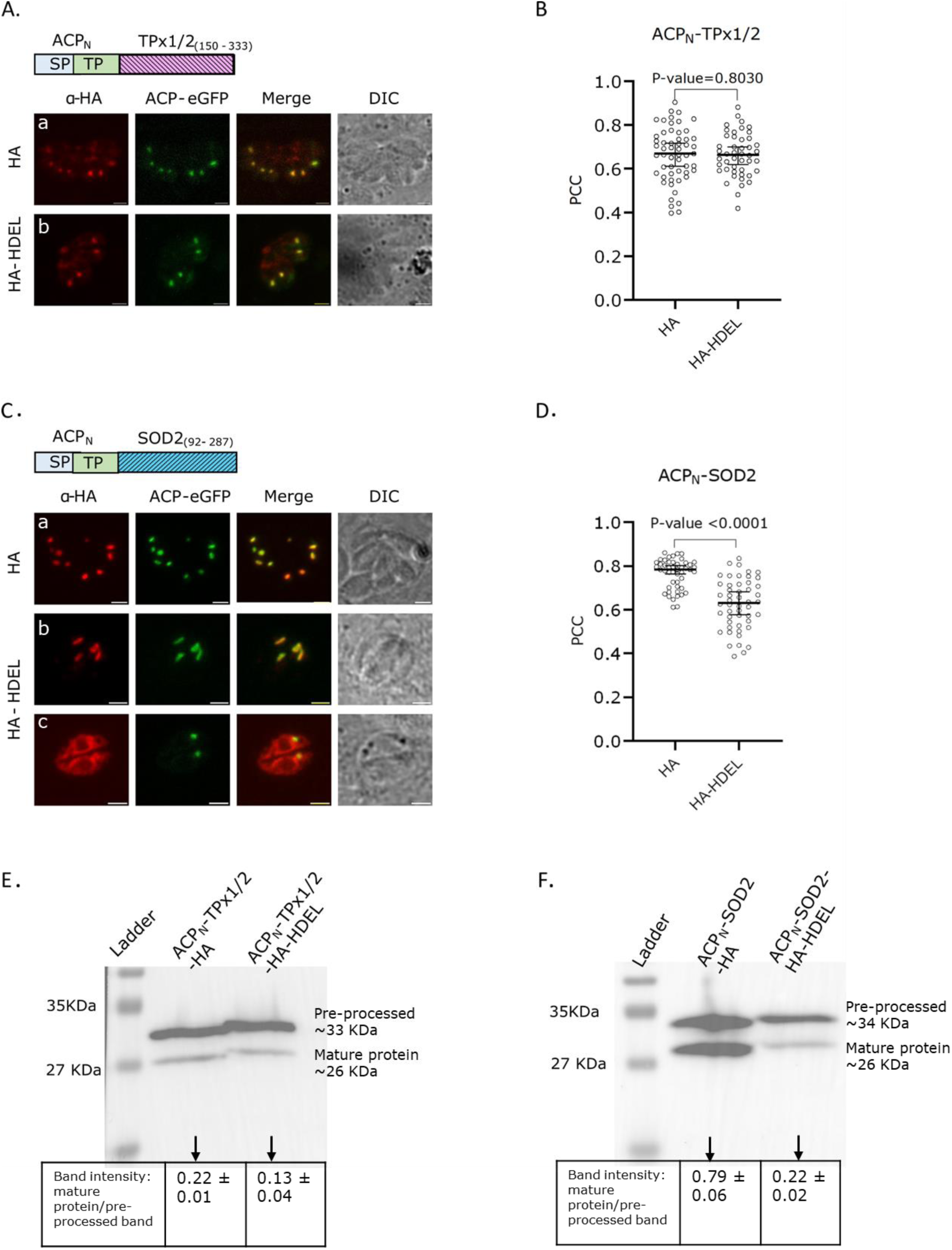
Analysis of the role of coding sequences of dually localized proteins in the trafficking route to the apicoplast. Microscopic images of parasites expressing tagged protein with or without HDEL motif are presented in red. The apicoplast marker is shown in green. **A)** *Tg*ACP_N_-TPx1/2, with apicoplast marker *Tg*ACP-eGFP (panel a and b) **B)** PCC values of *Tg*ACP_N_-TPx1/2 constructs with and without HDEL motif of *Tg*TPx1/2_N_-ACP showing median PCC value with 95% confidence interval with *Tg*ACP-eGFP **C)** Microscopic images of *Tg*ACP_N_-SOD2 with and without HDEL motif with apicoplast marker *Tg*ACP_N_-GFP, where panel a shows only apicoplast localization of *Tg*ACP_N_-SOD2-HA. In comparison, panel b and c shows two phenotypes of *Tg*ACP_N_-SOD2-HA-HDEL: only apicoplast and apicoplast + perinuclear staining construct, respectively **D)** PCC values of HA and HDEL constructs of *Tg*ACP_N_-SOD2 showing median PCC value with 95% confidence interval with the apicoplast marker *Tg*ACP-eGFP **E)** Western blot of *Tg*ACP_N_-TPx1/2 with and without HDEL sequence showing pre-processed protein of ∼33 kDa and the mature protein of ∼26 kDa. **F)** Western blot of *Tg*ACP_N_-SOD2 with and without HDEL sequence showing pre-processed protein of ∼34 kDa and the mature protein of ∼26 kDa. The quantitative analysis is based on data from 2 experiments. Scale bar = 2µm.

We used an alternate strategy to confirm these results. *Tg*ACP is an apicoplast luminal protein synthesized in a pre-processed form and processed during transit through the ER and the apicoplast membranes (Waller *et al*., 1998). Therefore, when apicoplast localization is compromised, *e.g.,* upon the addition of an HDEL sequence to a protein that uses a Golgi-dependent pathway, the ratio of pre-processed protein to mature protein will increase compared to that seen for the protein with no HDEL sequence. Conversely, if the protein takes a Golgi-independent route, there will be no change in the ratio of pre-processed to mature protein upon addition of an HDEL sequence, as apicoplast localization will not be compromised. Hence, we performed a western blot of *Tg*ACP_N_-TPx1/2-HA with and without the HDEL sequence. The western blot showed ratios of band intensity of 0.22 ± 0.01 and 0.13 ± 0.04 (mature protein to the pre-processed protein) for the protein with and without HDEL sequence, respectively (Fig. 4E). Therefore, there is no significant difference between the processing events in the two populations. This result further confirms a Golgi-independent route of *Tg*ACP_N_-TPx1/2-HA.

We next tested the ability of the *Tg*ACP signal sequence to drive the coding sequences of another dually localized protein, *Tg*SOD2 (92-287 amino acids), to the apicoplast. Parasites expressing *Tg*ACP_N_-SOD2-HA showed apicoplast localization in ∼60% of the population, and ∼40% of the population showed apicoplast localization with perinuclear staining (Fig. 4C, panel a). In contrast to this, parasites expressing *Tg*ACP_N_-SOD2-HA-HDEL showed apicoplast localization in ∼30% of the population and apicoplast localization with perinuclear staining in ∼70% of the population (Fig. 4C, panel b and c). The PCC analysis of *Tg*ACP_N_-SOD2 with and without HDEL was performed, and a significant decrease of the median PCC value of the parasites transiently expressing *Tg*ACP_N_-SOD2-HA-HDEL with a P-value <0.0001 was observed, indicating a Golgi-dependent route (Fig. 4D).

To validate the above findings, we performed western blots of parasite lysates expressing *Tg*ACP_N_-SOD2-HA, with or without the HDEL sequence. We observed that the ratios of band intensity of the mature protein to the pre-processed protein were 0.79 ± 0.07 in case of the protein without an HDEL sequence and 0.22 ± 0.02 for the protein with the HDEL sequence (Fig. 4F). This reduction in the level of the mature protein was consistent with the PCC values seen in Fig. 4D, and indicated that as the addition of an HDEL sequence reduced apicoplast localization, *Tg*ACP_N_-SOD2-HA uses the Golgi-dependent route for trafficking to the apicoplast. Therefore, although *Tg*ACP uses a Golgi-independent route to reach the apicoplast, the N-terminal signal sequence of this protein drives *Tg*TPx1/2 coding sequences through the Golgi-independent route, and *Tg*SOD2 coding sequences through the Golgi-dependent route. These data strengthen our hypothesis that a combination of both the N-terminal signal sequence and the coding sequences of a protein determines the pathway taken to reach the apicoplast.

## Discussion

In *T. gondii,* proteins such as *Tg*ACP and *Tg*FNR are exclusively localized to the apicoplast. Their N-terminal signal sequences, when fused to fluorescent reporters, reliably target these reporters to the apicoplast; hence, these fusion proteins have been used extensively as organellar markers (He *et al*., 2001; Pino *et al*., 2007; Amberg-Johnson and Yeh, 2019). In this report, we ask whether similar results are seen for the N-terminal sequences of *Tg*TPx1/2, a protein dually targeted to the apicoplast and the mitochondrion (summarized in Fig. 5). We found that when eGFP is used as a reporter driven by the N-terminal signal sequences of *Tg*TPx1/2, instead of dual localization to both the mitochondrion and apicoplast, it showed only mitochondrion localization. Surprisingly, a charged residue mutation, R24A, in the N-terminus of *Tg*TPx1/2_1-150_-eGFP should have resulted in apicoplast targeting (Mastud and Patankar, 2019), yet was also unable to drive eGFP to the apicoplast (Fig. 2). As similar results were reported for *Tg*SOD2 (Pino *et al*., 2007), the N-terminal targeting sequences of at least two dually targeted proteins are different from those of exclusive apicoplast proteins.

**Figure 5:**
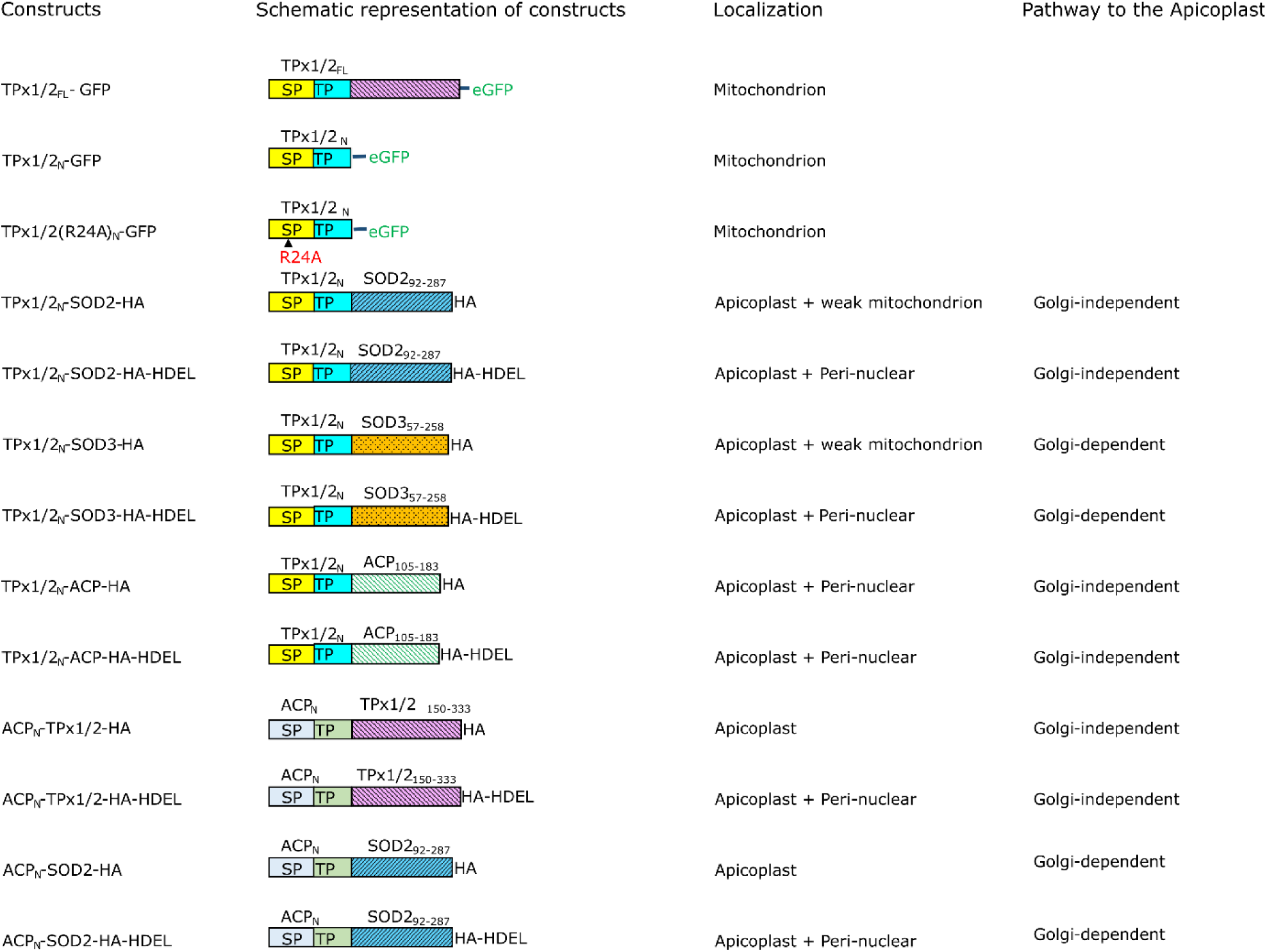
A summary of the localization and trafficking pathway of the chimeric proteins reported in this study. Schematic representation of chimeric proteins. The diagram represents the amino acids used to generate the chimeric proteins and the name assigned to each construct. All the chimeric proteins contain either eGFP or HA at the C-terminal to study their localization in the parasite. An HDEL motif was fused in-frame at the C-terminal end to study the trafficking pathway. SP-signal peptide, TP-transit peptide

*Tg*TPx1/2 contains ambiguous signals recognized by both the mitochondrial translocons as well as the signal recognition particle (SRP) for uptake in the ER, which is the first step in apicoplast trafficking. Earlier, a contest of strength or ‘tug-of-war’ was proposed between the two receptors for signal recognition of dually localized proteins with an ambiguous signal (Mastud and Patankar, 2019). After this early step, dually localized proteins use a Golgi-dependent pathway to reach the apicoplast. The presence of the eGFP reporter appears to interfere with one or more steps in this pathway to the apicoplast. Based on the complete lack of apicoplast or ER localization of eGFP fusions with the N-terminus of *Tg*TPx1/2, we propose that the very first step of apicoplast trafficking *i.e.* the targeting to the ER is being affected, allowing the mitochondrial translocons to win the tug-of-war. Two reports suggest mechanisms by which this might happen.

N-terminal signal peptides have similar properties in terms of positive charge, overall hydrophobicity, and a polar region (von Heijne, 1985). However, they greatly vary in their ability to translocate proteins into the ER lumen. For example, when non-native coding sequences are fused to the SP of proteins, there is a reduction in the translocation efficiency of the protein into the lumen of the ER (Kim *et al*., 2002). This report suggested that proteins are functionally matched with the N-terminal signal sequences for efficient translocation into the ER lumen. Proteins with inadequate translocation would be present in the cytosol (Dedhar, 1994; Holaska *et al*., 2001; Johnson *et al*., 2001), and for *Tg*TPx1/2, this would favor mitochondrion localization.

Another report that studied the plastid of the diatom *Phaeodactylum tricornutum*, showed that signals present in the coding sequences, especially the sequences found in the vicinity of the TP, can influence protein trafficking to their destination (Felsner, Sommer and Maier, 2010). Here, motifs that act as intrinsic TP sequences were uncovered in the eGFP protein when certain combinations of sequences were created during cloning. Similar findings were reported in higher plants where using chimeric proteins showed that merely adding a transit peptide to different coding sequences did not ensure correct chloroplast targeting (Lubben *et al*., 1989). Instead, correct targeting may be attributed to the additional signals present in the coding sequences or the protein’s secondary and tertiary structure (Wasmann *et al*., 1986; Lubben *et al*., 1989; Dabney-Smith *et al*., 1999). For example, the transit peptide of the small subunit of the Rubisco enzyme showed increased protein import efficiency in chloroplast after the inclusion of the first 23 amino acid residues of the mature protein when fused with coding sequences of different proteins (Wasmann *et al*., 1986).

Apicoplast localization was restored when eGFP was replaced with endogenous proteins, *Tg*SOD2 and *Tg*SOD3, as reporters (Fig. 3). These results suggest that dual organellar targeting of fusion proteins having the *Tg*TPx1/2(1-150) N-terminus is affected by both the ambiguous N-terminal signal sequence of *Tg*TPx1/2 and the coding sequences of proteins, which might contain signals that are missing in eGFP. A surprising result obtained with these proteins was their weak mitochondrial localization. Mitochondrial targeting sequences are found in the N-terminus of the protein and should not be affected by the coding sequences of the fusion protein. Again, one could invoke the unique feature of dually targeted proteins: competition between the receptors at the apicoplast and mitochondrion. *Tg*SOD2 and *Tg*SOD3 as reporters may tip this balance in favor of the SRP rather than mitochondrial translocons, as suggested by the two mechanisms described above.

What are the implications of fusion proteins showing different organellar localization? It has been proposed that evolutionarily, the acquired gene products from the endosymbiont do not attain signals for specific protein targeting to the new organelle immediately. Instead, these proteins explore different organellar environments before choosing the compartment where the proteins can form a complete biochemical pathway (Martin, 2010). For these expeditions, it would be favorable if the signal sequences and the coding sequences of proteins showed plasticity to be able to sample different organelles. It is tempting to speculate that our work provides a glimpse into the evolution of protein trafficking to the endosymbiotic organelles. The composite plasticity of the N-terminal and the coding sequences influences the final destination of the protein: the trafficking of *Tg*TPx1/2 fused with either eGFP or endogenous proteins to different compartments represents a molecular mechanism of exploring different destinations during the evolutionary process.

Proteins are trafficked to the apicoplast via two distinct pathways (reviewed in Mallo *et al*., 2018; Prasad, Mastud and Patankar, 2021); however, what factors determine which pathway to follow is still unclear. It was earlier suggested that the difference in the strength of the transit peptides present at the N-terminus of exclusive apicoplast proteins and the dually localized proteins could be the determining factor for the choice of the pathway (Prasad, Mastud and Patankar, 2021). This report shows, for the first time, that in addition to the N-terminus of a protein, the choice of trafficking pathway for apicoplast proteins is also dependent on the coding sequences, as summarized in Fig. 5. Further investigations are needed to decipher whether the coding sequences of proteins contains any additional motifs, or the trafficking information is present in the 3-dimensional structure, as seen for rice α-amylase (Kitajima *et al*., 2009).

In conclusion, in the field of apicoplast protein trafficking, N-terminal bipartite signal sequences have been considered both necessary and sufficient for trafficking to the destination. We show, for the first time, that the coding sequences of proteins have information, both for appropriate localization to the apicoplast and also for the choice of the trafficking pathway. This data opens up new areas of research in the field. For example, if the trafficking pathways to the apicoplast can be switched by making chimeric proteins, the functional importance of each of the pathways is of great interest. Further, searching for receptors in the ER and Golgi that recognize the N-terminal signal sequences and regions of the coding sequences of apicoplast proteins will shed light on the molecular players of apicoplast trafficking pathways. Finally, our results provide a word of caution to researchers in the field: fusion proteins using N-terminal sequences of apicoplast proteins must be tested empirically for their localization and trafficking pathways.

## Funding Source

We thank the Department of Biotechnology, Govt. of India, for research funding (Grant no. BT/PR13546/BRB/10/1423/2015) to SP and the PhD fellowship (DBT/2018/IITB/1149) to SA. We thank the Ministry of Education, Govt. of India and IIT Bombay for the PhD fellowships to AP and PM.

## Authorship contribution statement

SA, AP, PM and SP conceived and designed the experiments. SA, AP, PM performed the experiments. SA, AP and SP analyzed the data and prepared the manuscript. SA prepared the figures. SA, AP, PM and SP reviewed drafts of the manuscript.

## Declaration of competing interest

The authors declare that they have no known competing financial interests or personal relationships that could have appeared to influence the work reported in this paper.

## Acknowledgements

We acknowledge the Indian Institute of Technology Bombay for their Confocal Laser Scanning Microscope facility. We appreciate Manesh Joshi for constructing the pCTG-ACP_L_-HA plasmid and Geetanjali Mishra for her constructive feedback on the manuscript.

## Notes

### Competing Interest Statement

The authors have declared no competing interest.

### Summary of Updates

This manuscript version has been revised to update the representative images of the chimeric construct of *Tg*TPx1/2(R24A)N-eGFP, *Tg*TPx1/2-SOD2-HA, *Tg*TPx1/2-SOD3, and *Tg*ACP-SOD2. The Western Blot of *Tg*TPx1/2FL, *Tg*TPx1/2N-SOD2, and *Tg*TPx1/2-SOD3 in Figure 2 was also updated. Moreover, new Western blot data has been added for ACP-TPx1/2 and ACP-SOD2 with and without HA-HDEL motif at their C-terminal in Figure 4 to provide insights into the trafficking pathway of these constructs. As per the updated results, the model in Figure 5 is removed, and the text in the manuscript has been edited for better readability.

